# FEHAT: Efficient, Large scale and Automated Heartbeat Detection in Medaka Fish Embryos

**DOI:** 10.1101/2024.07.21.604466

**Authors:** Marcio Soares Ferreira, Sebastian Stricker, Tomas Fitzgerald, Jack Monahan, Fanny Defranoux, Philip Watson, Bettina Welz, Omar Hammouda, Joachim Wittbrodt, Ewan Birney

## Abstract

High resolution imaging of model organisms allows the quantification of important physiological measurements. In the case of fish with transparent embryos, these videos can visualise key physiological processes, such as heartbeat. High throughput systems can provide enough measurements for the robust investigation of developmental processes as well as the impact of system perturbations on physiological state. However, few analytical schemes have been designed to handle thousands of high-resolution videos without the need for some level of human intervention. We developed a software package, named FEHAT, to provide a fully automated solution for the analytics of large numbers of heart rate imaging datasets obtained from developing Medaka fish embryos in 96 well plate format imaged on an Acquifer machine. FEHAT uses image segmentation to define regions of the embryo showing changes in pixel intensity over time, followed by the classification of the most likely position of the heart and Fourier Transformations to estimate the heart rate. Here we describe some important features of the FEHAT software, showcasing its performance across a large set of medaka fish embryos and compare its performance to established, less automated solutions. FEHAT provides reliable heart rate estimates across a range of temperature-based perturbations and can be applied to tens of thousands of embryos without the need for any human intervention.

## Introduction

The heart is an orthologous organ system across all vertebrates (Monahan-Earley *et al*., 2013) and has its evolutionary roots deep in the metazoa, with conserved autonomous beating functionality ultimately controlled by orthologous transcription factors from arthropods to vertebrates (Souidi and Jagla 2021). As such, metrics describing the function of the heart, for example its heart rate, are beneficial physiological measures, long used in human physiology and widespread in the investigation of model organisms. The resting heart rate is influenced by both genetic and environmental factors. Small fish models, such as medaka (*Oryzias latipes*) (Aida, 1921) and zebrafish (*Danio rerio*) (Streisinger *et al*., 1981; Shin & Fishman, 2002; Lieschke & Currie, 2007), have been a source of discovery and functional characterisation due to the ease of husbandry and genetic manipulation. These models have enabled the discovery and validation of genetic variants, assessment of phenotypic drug safety pipelines, and toxicological studies, often using heart rate as an informative metric (Tonelli *et al*., 2020). The development and function of the fish heart are ancestral and comparable to those of the mammalian heart, mainly due to similar underlying molecular and cellular mechanisms that govern cardiac physiology and morphology. Both fish and mammalian hearts share conserved genetic pathways and structural proteins that play pivotal roles in heart development and function such as contraction, and response to stressors. See Jewhurst *et al*. (2016), Hu et al. (2020), and Nguyen et al. (2022) for comprehensive reviews. The rapid development of the cardiovascular system in fish allows its functionality to emerge in the relatively short time frame of approximately 48 hours in medaka post-fertilization (Gierten *et al*., 2020), and its small embryo size allows the generation of high throughput assays for embryo phenotyping (e.g., developing medaka fish embryos in a 96-well plate format).

It is obviously desirable to automate the measurement of heart rate from these embryos under different physiological conditions. A confounder to easy image analysis is the variable orientation of non-mounted embryos relative to the objective lens. As the heart is usually the only moving organ in the transparent zebrafish and medaka fish embryos, many orientations will provide a view of the heart movement (often the blood fluid), but these different views have different shapes and relative orientations of the heart. Several heart rate detection software packages for fish are available as open-source software (Gierten et al. 2020; Pylatiuk et al, 2014; Vedder et al, 2023). The software created by Gierten and collaborators is a peak counting method suitable for an autonomous analysis of a few samples at a time, as it requires human interaction as a pre-processing step often to select the region of interest for the assessment of the heart rate. The software created by Vedder and collaborators also requires some human interaction to select the heart region from a fluorescently labelled zebrafish heart. This method then offers two options for assessing heart rate: the user can fit a sine function to the variability of pixel intensities or apply a Fourier transform (FT) to the data. These two examples require either fluorescent labelling of the heart or manual selection of the heart region, or both, which means that applying them to hundreds or thousands of fish can be laborious. The third example is the software created by Pylatiuk and collaborators, which can automatically detect the heart using a change-in-color technique applied to each pixel in the image. The Gierten and Pylatiuk software is written in MATLAB, which is entirely appropriate for small scale, manual operation but less easy to deploy at scale.

To facilitate the automation of reliable and efficient quantification of heart rates at a large scale, we have developed an open source (Python) and user-friendly application, FEHAT - the Fish Embryo Heartbeat Assessment Tool. This software package automatically selects a region of interest, regardless of orientation, and then applies a FT function to accurately characterise the major frequency components of fish embryos over time. The input to the FEHAT tool is multiple videos, often obtained from imaging a 96-well plate, and the output is a single results file containing the heart rate frequency for each embryo, as well as some other informative derived information about the signal. No human interaction is necessary, as the embryo and heart region detection, frequency transformations, and quality control are all done automatically. This allows very high levels of scalability.

This paper describes the main image processing and analysis of the FEHAT software and presents its concordance with established, less automated software. We also explore the use of other information present in the image and discuss the practical steps for large-scale automation. In our hands, we have successfully run this software over 10,000 videos in a fully automated manner. We provide FEHAT as an open-source Python-based software package (https://github.com/birneylab/FEHAT).

## Methods

To develop and validate the software we used a variety of medaka fish strains available at Heidelberg University, where the fish stocks are kept in a closed system. The husbandry of medaka was conducted in compliance with local animal welfare regulations (Tierschutzgesetz §11, Abs. 1, Nr. 1) and European Union guidelines with a permit number of 35–9185.64/BH Wittbrodt, Regierungspräsidium Karlsruhe. They were raised and maintained following previous protocols. All the analysis was done at the embryo stage. The local representative of the animal welfare agency oversees the fish facility.

We used five distinct data groups to validate our software and ensure accuracy, which varied in genetic background, temperature, frame rate, and operator. In this paper, we have named these Datasets 1, 2, 3, 4, and Cabs for convenience. Dataset 1 represents offspring of a cross between two inbred strains of fish, whereas Dataset 2 and Dataset 3 denote a combination of mock injected and crispant fish available in the facility. Dataset 4 is composed of data from different strains selected randomly. Finally, dataset Cabs are a specific strain often used as the standard laboratory control. We processed Datasets 1 to 4 using previously developed semi-manual, peak counting based software (Gierten et al., 2020), which provides a high accuracy measurement of heartbeat but requires a manual step to identify the region of interest in all embryos in the test plates. These semi-manual datasets and our automated FT calls are available as **supplementary material**. The value recorded for videos without a visible heart is named “no peak counting available” or “no FT calling available”. The heart rate in fish shifts with changing temperatures. To explore the software applicability on a spectrum of heart rates ranging from 70 beats per minute (bpm) to 286 bpm, measurements from three different temperatures were assessed. The number of data points and temperature varied across datasets: 925 for Dataset 1 across 21°C and 28°C, 96 for Dataset 2 at 21°C, 288 for Dataset 3 across 21°C, 28°C, and 35°C and 96 for Dataset 4 at 35°C. These datasets were used to compare the semi-manual peak counting approach against our automated FEHAT software. For Cabs, 1296 embryos were sampled at 21°C, 28°C, and 35°C, and embryos were used to compare two subsequent measurements (loops) using the FEHAT software only.

The workflow of the heartbeat extraction using a FT in transparent medaka fish embryos is illustrated in **Figure 1** and explained in the results section.

**Figure 1:**
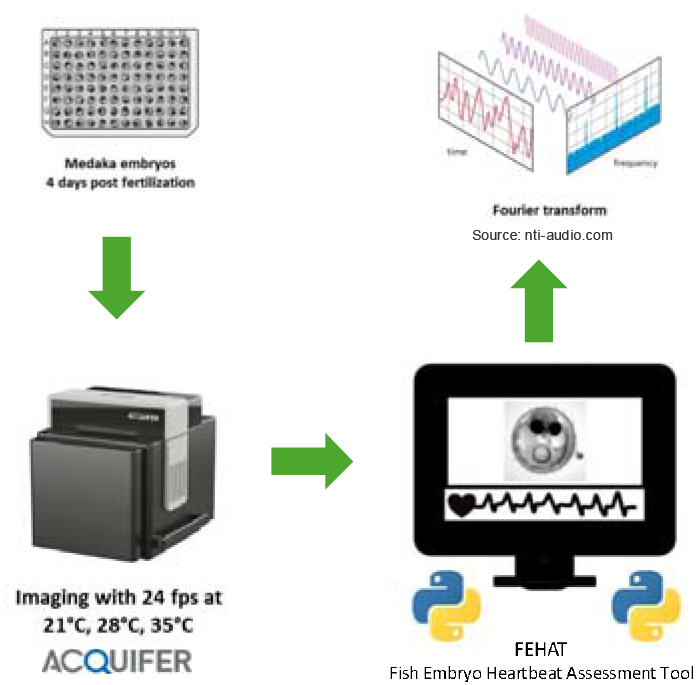
Flow diagram showing the main steps to measure heartbeat using transparent Medaka fish Embryos on the Acquifer imaging device and FEHAT.

### Automated microscopy

The 96-well microtiter plates containing medaka embryos at stage 32, four days post fertilisation (dpf) were exposed to specific temperatures (21, 28, and 35°C) using a temperature-controlled incubation lid (LUXENDO GmbH, Heidelberg, Germany). These plates were automatically imaged using an ACQUIFER Imaging Machine, a wide-field high content screening microscope equipped with a white LED array for bright-field imaging, an LED fluorescence excitation light source, an sCMOS (2048□×□2048 pixel) camera, a stationary plate holder, and movable optics. The bright-field channel detected the focal plane using an autofocus software algorithm. Images were captured as a 10-second movie with either 13 or 24 frames per second, all at the same z-axis position determined by the built-in software autofocus. Further details about this setup can be found in (Gierten et al., 2020). The resulting output from capturing an entire 96-well plate was a total of 12,480 or 23,040 frames depending on the framerate, (130 or 240 frames for each embryo, respectively), saved in a TIFF format.

### Software outline

Once the data has been acquired, the following technical steps are carried out:

The first step involves detecting the embryo in the images and cropping if needed, avoiding unnecessary pixels to be analysed. This both greatly reduces memory and compute requirements and importantly does not require human interaction during the cropping. After converting the images to grayscale, the software performs edge detection in each frame, followed by a Hough Circle Detection to obtain the exact region of the embryo. The embryo’s location is considered the most common coordinates in the set of frames assessed, and the images are cropped around the maximum area around the embryo edges. Importantly, the process only continues to the next step if less than 5% of frames are empty, as the camera used for image acquisition can occasionally generate corrupted videos, resulting in an extensive sequence of black frames and invalidating the sample. All steps regarding image manipulation have been done using OpenCV (2022) and NumPy (2023).

The second step is to normalise across frames. Typically, the images are acquired with variable lighting, presenting a significant variation in pixels’ brightness value across frames. We perform a linear normalisation to correct for these variances.

The third step automatically selects the heart region of interest (see the full workflow for this step in **Figure 2**). To extract only the heart region from the set of images, FEHAT calculates the absolute rolling difference with a defined window size (default of 2 frames) after blurring them. Then, triangle thresholding is applied to the two frames’ absolute difference. This produces a mask with regions of more significant variations in pixel intensities over the window. Any resulting noise is removed using morphological opening, maintaining only areas of substantial size. We call these initial regions of interest (IROI, **Figure 2** “Pixels emitting change”).

**Figure 2:**
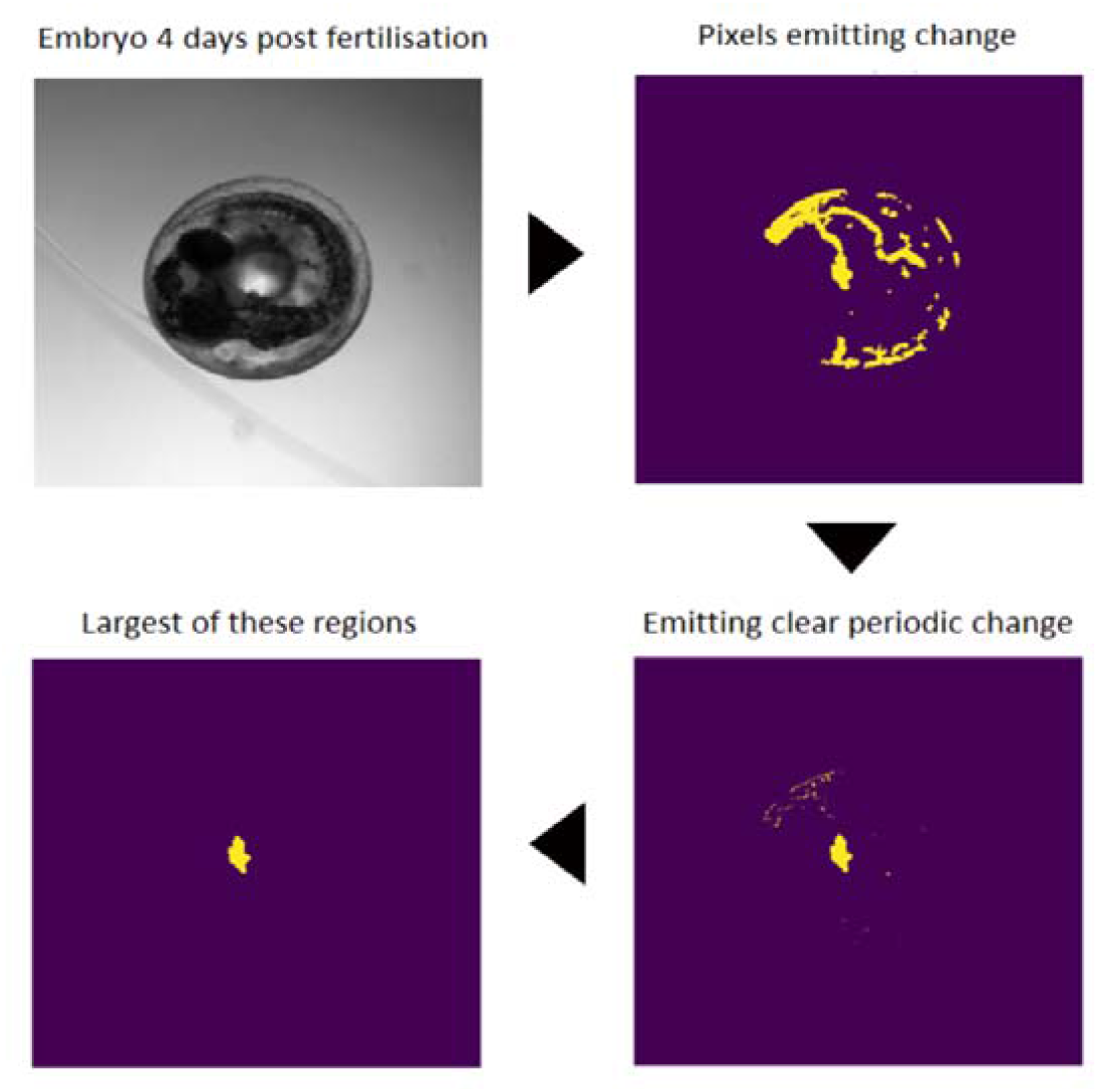
The four main steps to get the right heart region. The videos are obtained from an Aquifer machine (upper left); only pixels with periodic changes are selected (upper right); a FT is applied (bottom right); the region with the biggest area is selected as the heart region (bottom left).

In the fourth step, IROIs are filtered based on clear periodic pixel changes and object sizes. The aim here is to identify, among all IROIs, the one that is most likely to be the heart. Firstly, using the mask created in the previous step, which selected only pixels with variation above a certain threshold, FT is applied to obtain the frequency amplitude in each pixel. Regular and periodic movement, such as the heart, will have a large FT peak at a specific frequency, whereas less regular movement, such as blood flow in the external veins of the embryo will not have a large FT peak. We then use a user-defined range of acceptable FT peaks (for medaka fish hearts, between 0.5 and 5 Hertz). Finally, we calculate a signal-to-noise ratio (SNR) of the top FT peak to all others and select pixels within an acceptable user defined SNR range. A new mask is created from these periodic pixels, and the largest periodic region is selected. As discussed below, an important aspect of using the FT analysis is that when the field of view encompasses the two chambers of the heart both chambers will beat at a similar frequency, but with different phases.

In the fifth step, the signal in the heart region is calculated using the mask described above, and the heart rate is estimated using the FT from these pixels. To obtain a single heart rate value from a set of pixels, kernel density estimation is used to estimate the most common frequency component, which can then be used to estimate the heart rate.

**Figures 3-a** and **b** represent the kernel density estimation of the final output of the FT analyses applied to a set of pixels presented in the ROI. In **Figure 3a**, the height of the lines in the y-axis represents the frequency amplitude in each pixel (z-axis) in each frequency (x-axis). **Figure 3b** is an “upper” vision of **Figure 3a**, with white intensities representing the height of the peaks (the frequency amplitude). The FT reports naturally in Hz (beats per second), and we multiply by 60 to get the more biologically expected beats per minute (bpm).

**Figure 3:**
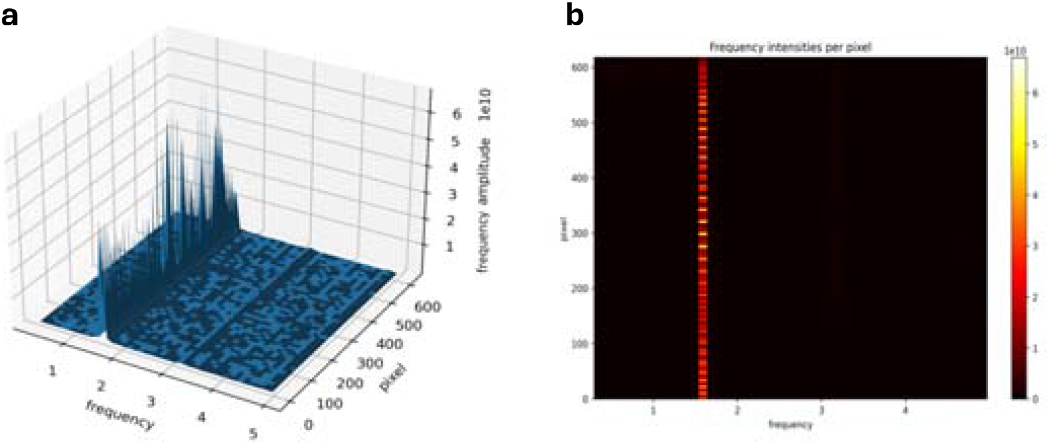
Example of the FT spectrum. (a) 3D representation of the pixels in each frequency bin as calculated by FT. The chosen frequency for the heart rate estimate is often the highest for each pixel, and (b) is the same as above but using a heatmap representation—the highest pixels for the brighter regions in the picture.

### Data cleaning post automation

We developed robust quality control procedures that can be automatically applied after the main extraction of heart rate to improve data reliability.

Often, slight changes in orientation over the image capture time will provide different views of the heart; in some orientations, such as an axial view, the two-chambered heart can give a robust readout of twice the heart rate. Often, even a slightly alternative orientation provides a clearer view of each chamber. When this automated heart rate measurement is deployed in two separate technical measurements, a common error mode is that one measurement is double the frequency of the other. To exploit this, we developed a procedure that when multiple measurements are made on the same fish embryo (technical replication) to detect pairs of measurements where one measurement is twice the other, and the other measure falls within a group average. The “double measurement” is halved in these cases, and both are averaged. Alternatively, an average is taken if both measurements fall within the expected range. This automatically and robustly handles the system’s most common source of variation - the detection of the second harmonic.

We also discard measurements which are below specific limits (default is 70 bpm), but this can be changed by the user and is specific to each experiment (eg, chemical or drug treatment).

## Application to diverse medaka datasets

We applied this method to four real datasets representing a range of scenarios from medaka fish embryo investigations.

**Figure 4a** shows the result of running FEHAT on embryos from the Cab dataset through two time-separated assessments, where each time window is analysed separately as Loop1 or Loop2. Most cases have very similar measurements, but a subset shows cases where one loop is twice the frequency of the other, suggesting an embryo movement between measurements with an axial view of the heart (see above). One limitation of the FT approach is that the necessary discrete nature of the FT leads to a discrete read out of the frequency, shown in the **Figure 4** and **Figure 5** by the grouping of the points.

**Figure 4:**
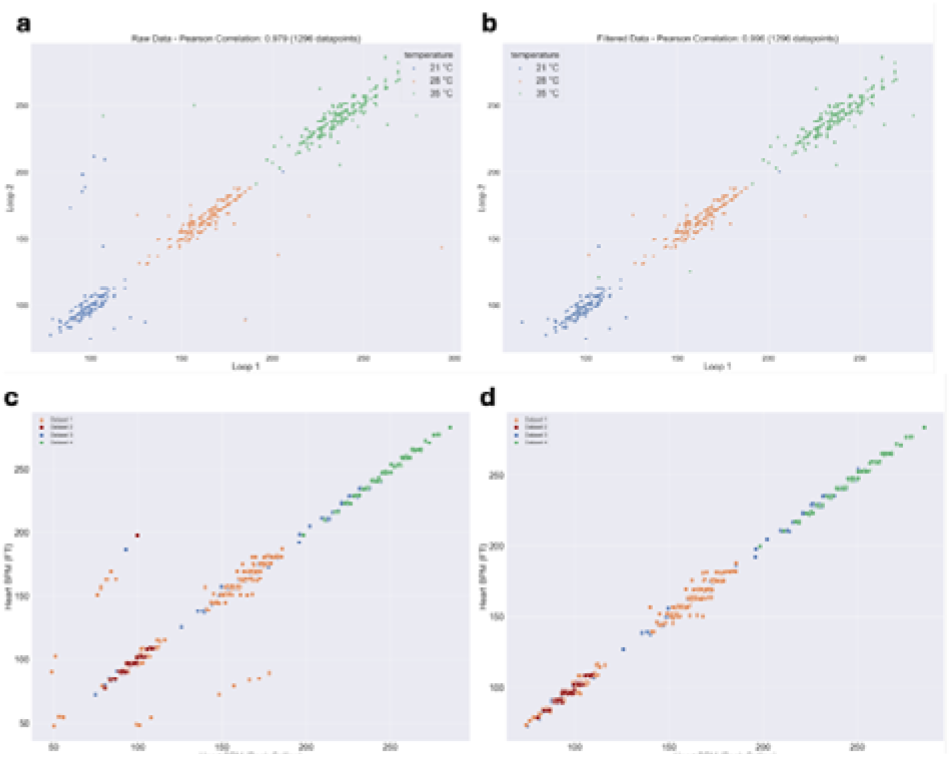
Loop 1 versus loop 2 at 3 different temperatures. (a) Heart rate estimate using FT before correction of the double harmonics. (b) Heart rate estimate using FT after correction of the double harmonics. (c) Peak calling approach versus FT across 4 different datasets prior to double harmonic correction. (d) Peak calling approach versus FT across 4 different datasets after double harmonic correction.

**Figure 5:**
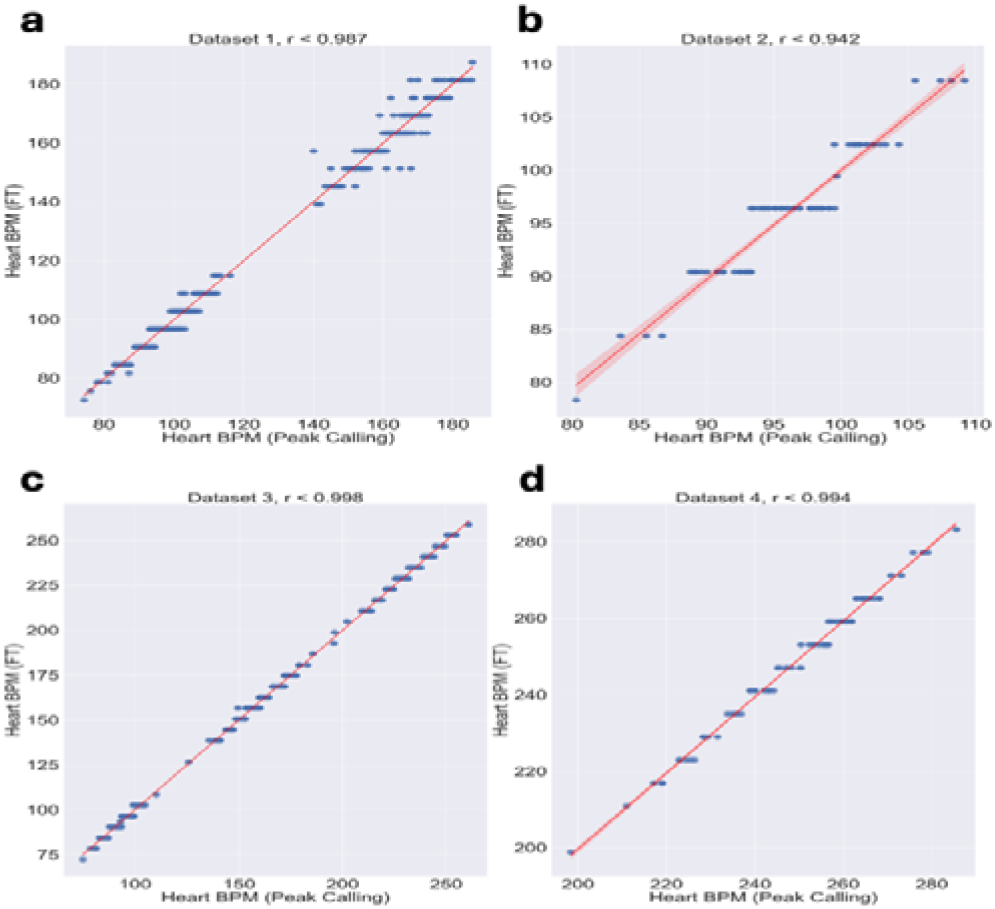
Scatterplots of all four datasets with their respective value of “r” of Pearson after the cleaning and transformation procedure.

We also compared the results of our fully automated system to a previous manual selection of pixels followed by a peak counting assessment of frequency and reassuringly found a high level of concordance between the two methods (**Figure 4c and 4d**). **Figure 5a-d** shows the same four datasets run through the semi-manual procedure (peak finding) and our new automated procedure. The automated results are highly correlated with the previous measurement (r values: 0.981, 0.911, 0.997, and 0.99) showing a very consistent heartbeat measurement. This also shows that the discrete nature of the FT transform does not lead to a large change in variance compared to a peak finding approach.

One of the benefits of using the FT scheme is that the frequency and the phase of the beat are separated, the phase being the offset of the beat relative to the start of the video, measured in radians. We explored the distribution of phases in embryos, clustering pixels by the same phase (**Figure 6**).

**Figure 6:**
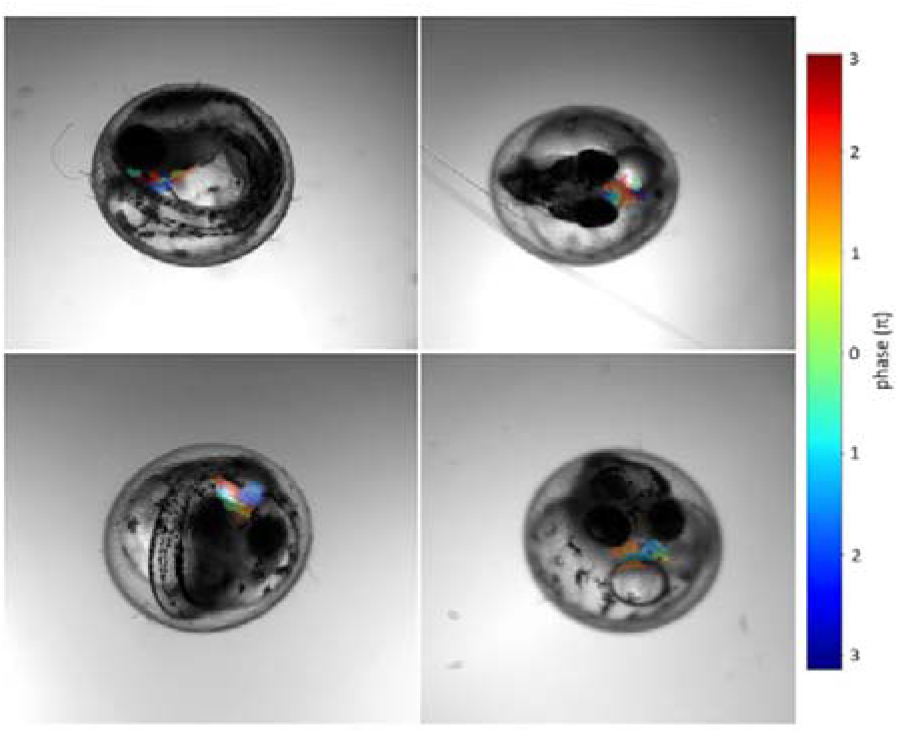
Colormap of phases value returned by the FT on the embryo ROI (region of interest).

In cases with more than 2000-3000 pixels in the fully connected region, there were often three main clusters of phases. The two more extreme phases were often counter beating, with an offset of about π radians; this offset implies two regions which are beating out of phase. The intermediary phase is probably an interaction between the movements of the two chambers. By visual inspection, when we have a ∼ π phase difference, these are cases where both the atrium and the ventricle are visible, and it shows that the FT can provide more information than just the main modal frequency.

In addition to the output of the main frequency of the heart rate, FEHAT also outputs other measurements associated with the automated region capture and FT features. These features have been useful for ranking images for quality control, quality assurance, and exploring datasets. For example, a common failure mode is due to movements of the embryo during the video acquisitions and if it occurs before 5 seconds the resulting frequency profile will be too short for an accurate analysis.

## Discussion

In this paper, we have developed robust software that can automatically determine the heart rate of medaka fish embryos in nearly any orientation. The software has been compared to an alternative approach that combines the manual selection of the most important pixels with a peak counting approach. Our software has an entirely automatic selection of key pixels and then uses a discrete FT to derive frequencies. A particular focus of this software is handling the diversity of possible orientations that developing fish embryos (in our case, medaka) have in a 96-well plate format. We have automated some key post-calling quality control checks, including handling the most common error: when the heart’s orientation gives rise to a double heartbeat (twice the frequency) in the setting of two independent orientations of the same embryo. In practice, even small orientation changes will rectify the double beat phenomena, meaning that sequential measurement with a small time gap provides a robust heart rate measurement. Both methods are highly concordant over multiple genetic backgrounds and temperatures, and our automated method can be scaled on modern compute farms to process large numbers of videos robustly.

We choose to use the FT to characterise the heart rate. The major benefit of the FT is that two chambers of the heart will be beating at the same frequency but with different phase offsets. The FT automatically resolves this issue, meaning our software does not have to distinguish these separate compartments. In addition, the FT provides a richer definition of the dynamics of the heart by being able to see other frequencies in the heart rate function and explore the phase offsets between regions of the heart. Clearly, this data will be more susceptible to orientation effects, but it provides a richer numerical phenotype from these embryos than a single measurement. In systematic studies, we are currently exploring whether these richer measurements correlate with biological features (e.g., genetics or perturbants). One consideration of using the FT is that the resolution on the frequency of heartbeats is directly related to the length of the videos, a feature that can be seen by the discrete distribution of the data in the scatterplots shown.

We have now used this software in two large-scale campaigns (analysis ongoing), and the method is robust to 10,000s of embryos measured by different versions of the Acquifer machine (e.g., frame rate) and by different operators. In our hands, using a Load Sharing Facility with hundreds of compute hosts available, 24,000 samples can be processed within a day, though much of the complication is in storing, transferring, and accessing the video data.

Our software, FEHAT is fully open source (https://github.com/birneylab/FEHAT), so interested parties can download and adapt the software to other settings. The software system can ingest videos in 96-well plate format, such as from the ACQUIFER machine, but the fundamental core of the method is likely to apply to many scenarios. We expect this software can be easily adapted to other transparent embryo heartbeat scenarios, including zebrafish, but of course it will need customisation for each organism and experimental set up. We have not ported the software ourselves as we do not have access to large scale zebrafish or other datasets, but we look forward to a broader community using this software and welcome contributions to the code base.

## Conclusion

Our software was built to automate the extraction of the heart rate of thousands of fish embryos quickly and easily, without the necessity of any human interaction to identify the embryo nor the region of interest. We have used this software successfully in 10,000s of embryos with no human intervention in the individual embryo heart rate determination. By making it accessible as an open-source project, we hope to contribute significantly to developing research in heartbeat quantification in fish and other animals.

